# Gene Information eXtension (GIX): effortless retrieval of gene product information on any website

**DOI:** 10.1101/532523

**Authors:** James D.R. Knight, Payman Samavarchi-Tehrani, Mike Tyers, Anne-Claude Gingras

## Abstract

Retrieving information about genes and gene products is a constant and time-consuming aspect of systems biology research. While there are many high quality and well-designed resources to fulfill this need, they require the user to navigate to different and often complex websites, execute a search, select the desired result and then view retrieved information. This task can be a repetitive, burdensome and disruptive process, for example when exploring the results of large scale genomics or proteomics screens or reading an online article. To address this issue we have developed a browser extension for Google Chrome and Mozilla Firefox called GIX, for Gene Information eXtension, that allows users to retrieve customizable gene product information - especially as it relates to proteins and their expression and functions - directly on a website without having to navigate to another page.

---

Retrieving information about genes and gene products is a constant and time-consuming aspect of systems biology research. While there are many high quality and well-designed resources to fulfill this need, they require the user to navigate to different and often complex websites, execute a search, select the desired result and then view retrieved information. This task can be a repetitive, burdensome and disruptive process, for example when exploring the results of large scale genomics or proteomics screens or reading an online article.

To address this issue we have developed a browser extension for Google Chrome and Mozilla Firefox called GIX, for Gene Information eXtension. GIX allows users to retrieve customizable gene product information – especially as it relates to proteins and their expression and functions – directly on a website without having to navigate to another page. Simply double-clicking (or alternatively, mouse dragging) on a gene name or supported accession number (Ensembl, Entrez, neXtProt, RefSeq or UniProt) will open an information panel on the current page (Fig. 1). This panel includes gene synonyms, the full gene name, alternative names, the size and molecular weight of its canonical protein product, the UniProt description, protein domains and regions, GO terms, protein localization, RNA tissue expression, associated diseases or phenotypes, pathways, protein interactors (from BioGRID and IntAct) and links to external resources (Ensembl, NCBI and Uniprot for all species, and organism-specific databases: dictyBase, FlyBase, MGI, neXtProt, PomBase, SGD, TAIR, WormBase, Xenbase and ZFIN). Gene Info also offers an alternative tooltip mode that simply provides links to these external resources. The extension is fully customizable, allowing the user to control the information they see, and supports queries for *Homo sapiens* and ten model organisms: *Arabidopsis thaliana, Caenorhabditis elegans, Danio rerio, Dictyostelium discoideum, Drosophila melanogaster, Gallus gallus, Mus musculus, Saccharomyces cerevisiae, Schizosaccharomyces pombe* and *Xenopus laevis*. The extension provides a search bar for entering queries manually and an online “workspace” for pasting gene lists from desktop applications for quick queries with GIX. While double-clicking to retrieve results is not possible on websites with embedded content such as Google Docs or PDFs, querying with the search bar does work on such webpages.

**Figure 1.**
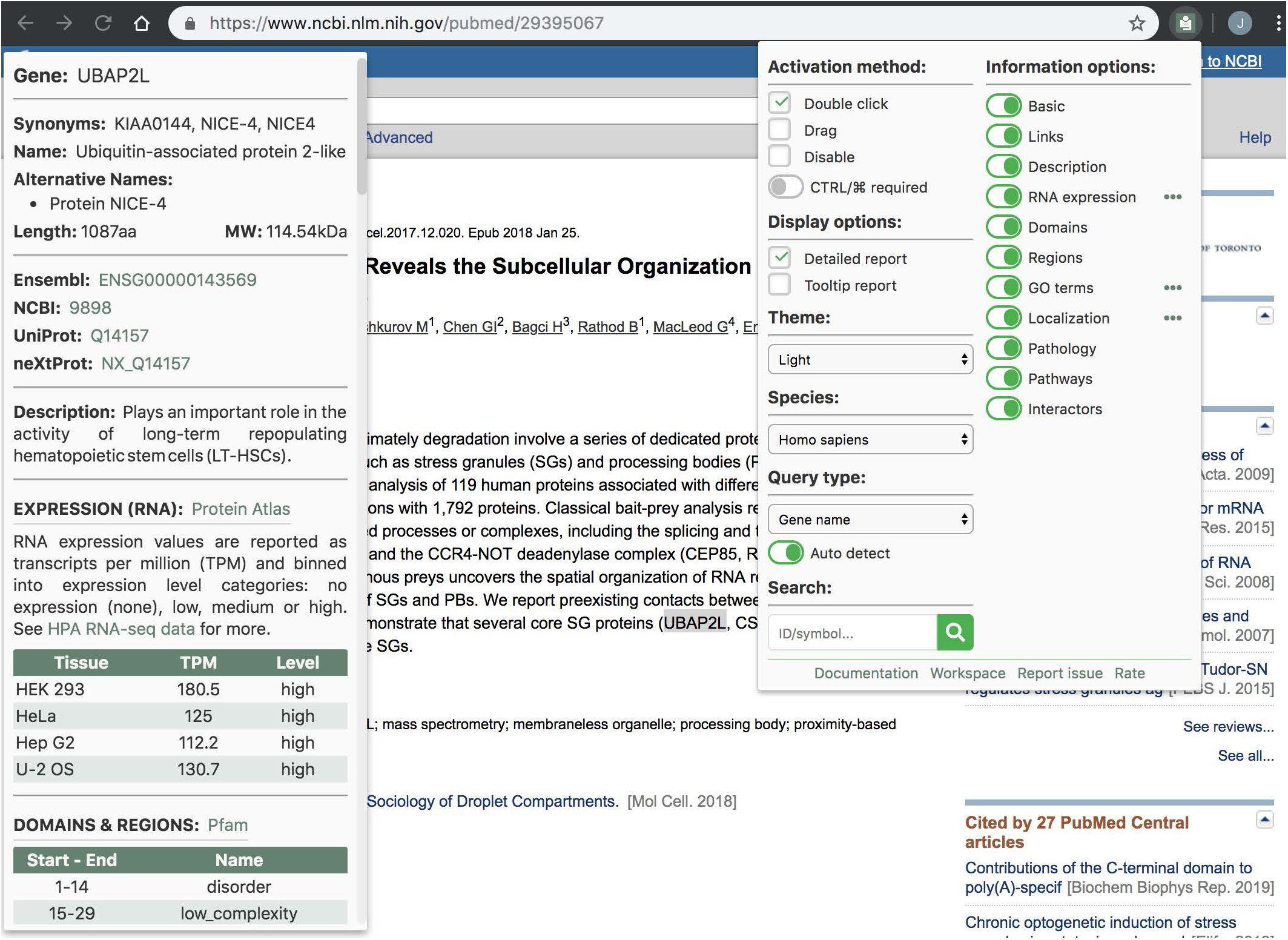
Screenshot of GIX in the Chrome browser. Double-clicking on a gene name (in this case UBAP2L) opens an information panel (left side) displaying information about the query. The extension has a number of settings that can be customized by clicking on its icon in the browser toolbar (right side).

GIX collates data from BioGRID (https://thebiogrid.org^1^), Compartments (https://compartments.jensenlab.org^2^), GO (http://www.geneontology.org^3, 4^), HUGO Gene Nomenclature Committee (https://www.genenames.org^5^), Human Protein Atlas (https://www.proteinatlas.org^6, 7^), IntAct (https://www.ebi.ac.uk/intact^8^), OMIM (https://www.omim.org^9^), Pfam (https://pfam.xfam.org^10^), Reactome (https://reactome.org/)^11^ and UniProt (https://www.uniprot.org^12^). The GIX database is updated monthly to incorporate changes from these resources. GIX is available for free without restriction at the Chrome Web Store and the Firefox Add-on site. Download links, documentation, a tutorial video and source code can be found at https://gene-info.org.

## ACKNOWLEDGEMENTS

We are grateful to all members of the Gingras lab for feedback on the extension. We acknowledge funding from the Governments of Canada and Ontario through Genome Canada, Ontario Genomics and the Ontario Research Fund (OGI-139 to A.-C.G. and RE08-065 to A.-C.G.), the Canadian Institutes of Health Research (Foundation grant FDN143301 to A.-C.G) and the National Institutes of Health Office of Research Infrastructure Programs (R01OD010929 to M.T. and K. Dolinski). This research was enabled in part by support provided by Compute Canada (www.computecanada.ca). A.-C.G. is the Canada Research Chair in Functional Proteomics and the Lea Reichmann Chair in Cancer Proteomics; M.T. is the Canada Research Chair in Systems and Synthetic Biology.

## AUTHOR CONTRIBUTIONS

J.D.R.K., P.S.T. and A.-C.G. conceived of the extension. J.D.R.K. wrote the code. M.T. provided input on the extension. J.D.R.K. and A.-C.G. wrote the manuscript with input from P.S.T. and M.T.

